# Distinct patterns of cortical manifold expansion and contraction underlie human sensorimotor adaptation

**DOI:** 10.1101/2022.06.09.495516

**Authors:** Daniel J. Gale, Corson N. Areshenkoff, Dominic P. Standage, Joseph Y. Nashed, Ross D. Markello, J. Randall Flanagan, Jonathan Smallwood, Jason P. Gallivan

## Abstract

Sensorimotor learning is a dynamic, systems-level process that involves the combined action of multiple neural systems distributed across the brain. Although we understand a great deal about the specialized cortical systems that support specific components of action (such as reaching), we know less about how cortical systems function in a coordinated manner to facilitate adaptive behaviour. To address this gap in knowledge, our study measured human brain activity using functional MRI (fMRI) while participants performed a classic sensorimotor adaptation task, and used a manifold learning approach to describe how behavioural changes during adaptation relate to changes in the landscape of cortical activity. During early adaptation, we found that areas in parietal and premotor cortex exhibited significant contraction along the cortical manifold, which was associated with their increased covariance with regions in higher-order association cortex, including both the default mode and fronto-parietal networks. By contrast, during late adaptation, when visuomotor errors had been largely reduced, we observed a significant expansion of visual cortex along the cortical manifold, which was associated with its reduced covariance with association cortex and its increased intraconnectivity. Lastly, we found that individuals who learned more rapidly exhibited greater covariance between regions in the sensorimotor and association cortices during early adaptation. Together, these findings are consistent with a view that sensorimotor adaptation depends on changes in the integration and segregation of neural activity across more specialized regions of unimodal cortex with regions in association cortex implicated in higher-order processes. More generally, they lend support to an emerging line of evidence implicating regions of the default mode network in task-based performance.

---

Adaptive behaviour depends on aligning one’s actions with the external constraints present in a given situation, and updating this mapping in response to new demands [1, 2]. Contemporary perspectives on brain function suggest that this process relies on cooperation between brain systems specialized for implementing behaviour in the moment, and those that help adapt behaviour in the face of a changing environment. In the field of motor learning, much focus has been placed on identifying the contributions of several sensorimotor brain areas whose activity varies over the course sensorimotor adaptation, a key form of learning by which the brain adjusts movement through trial-and-error [3–9]. For instance, it is well-understood that adaptation is supported, in part, by an implicit learning process wherein discrepancies between expected-versus-actual sensory outcomes (i.e. sensory prediction errors) are computed within the cerebellum [6, 8, 10, 11]. These sensory prediction errors serve as a ‘teaching’ signal to recalibrate subsequent motor commands in cortical sensorimotor regions, such as parietal, premotor, and motor cortex [12, 13], and thus gradually reduce errors over time.

In addition to this cerebellar-dependent learning process, emerging evidence indicates that sensorimotor adaptation is supported by explicit learning processes that presumably involve brain regions in association cortex [14–16]. This explicit component to adaptation uses knowledge about the change in environmental parameters in order to generate deliberate (and strategic) compensatory movements to minimize movement errors [17–19]. Considerable behavioural evidence suggests that explicit and implicit learning processes operate in parallel [15, 20], and that a greater relative contribution of explicit learning during the initial phases of adaptation, when errors are largest, results in a rapid error reduction [18, 21, 22]. Consistent with this notion, there is strong evidence that faster adaptation across individuals results from a greater recruitment of explicit processes, as compared to individuals who adapt more slowly [19]. Explicit learning is considered to be a largely cerebellum-independent process, involving the recruitment of higher-order cortical areas that exert top-down control over the sensorimotor system [14, 15]. To date, neuroimaging and lesion studies have mainly implicated brain areas involved in cognitive control and working memory, such as dorsolateral prefrontal cortex and parietal cortex [15, 23, 24], as supporting explicit learning processes.

However, an intriguing possibility is that this cortical involvement also extends into constituent areas of the default mode network (DMN), a collection of distributed areas implicated in the upper echelon of higher-order, or transmodal, processing [25]. Medial regions of the DMN, such as medial frontal cortex and posterior cingulate cortex, have been implicated in general shifts in strategy [26, 27] and are increasingly recognized as supporting several aspects of task-based cognition [28–31], commensurate with the role of the DMN in internal mentation [32–34]. It has been hypothesized that the broad contribution of the DMN to cognition (including during tasks) can be accounted for by its functional interactions with unimodal regions within sensory and motor networks [35–37]. These interactions are thought to be enabled through its positioning on the cortical mantel, located equidistant between unimodal systems involved in perception and action, which would allow it to uniquely encode and integrate features of whole brain activity [35, 38]. Taken together, the recruitment of explicit learning processes during adaptation, which are cognitive and strategic in nature, is expected to engage a much broader cortical functional architecture beyond conventional sensorimotor cortical regions.

In order to explore and characterize the widespread involvement of cortex during sensorimotor adaptation, our study leverages advanced manifold learning approaches that provide a compact, low-dimensional description of changes in the overarching cortical functional architecture. Recent electrophysiological work has established that low-dimensional manifolds can provide a compact description of the covariance of neural population activity within regions of the premotor and motor cortices [39–41]. At the same time, studies in other domains have applied the same logic to establish that low-dimensional representations of cortical activity can be a useful description of the macroscale organization of neural activity [42, 43]. Recently, whole-brain manifolds, or gradients, have provided insight into the low-dimensional organization of brain structure and morphometry [44–46], intrinsic brain activity during rest [38], and changes in brain organization in clinical disorders [47–49] and throughout the lifespan [50–52]. Here, we applied this approach to gain insight into how distributed cortical activity is coordinated during sensori-motor adaptation, and how this changing cortical landscape unfolds across different phases of learning. Specifically, by estimating the relative positions of cortical brain regions in a connectivity-derived manifold space, and understanding how these change in response to an environmental perturbation, we aimed to capture the evolving landscape of brain activity that supports sensorimotor adaptation, as well as the features of this activity that relate to better or worse learning performance.

## Results

We had participants (*N* = 32) perform a classic visuomotor rotation task [53] during functional MRI (fMRI) scans, in which they launched a cursor from an initial centre position to a cued target that could be located in one of eight encircling positions on a visual display (Fig. 1A). Participants launched the cursor by applying a brief isometric directional force pulse on an MRI-compatible force sensor. Following a Baseline block (120 trials), in which the cursor direction directly matched the force direction (Fig. 1A, *top*), the cursor was rotated 45^◦^ clockwise relative to the force direction for a remaining 320 trials (Fig. 1A, *bottom*). Overall, we found that participants’ angular error was low during Baseline, indicating that individuals could perform the task with high accuracy (Fig. 1B). Following the onset of the rotation, errors increased significantly as participants had to learn how to counteract the cursor rotation to successfully hit each target (by aiming their hand in a 45^◦^ counterclockwise direction). Over time, participants were able to reduce their error to near-Baseline levels of performance, indicative of successful adaptation (Fig. 1B).

**Fig. 1.**
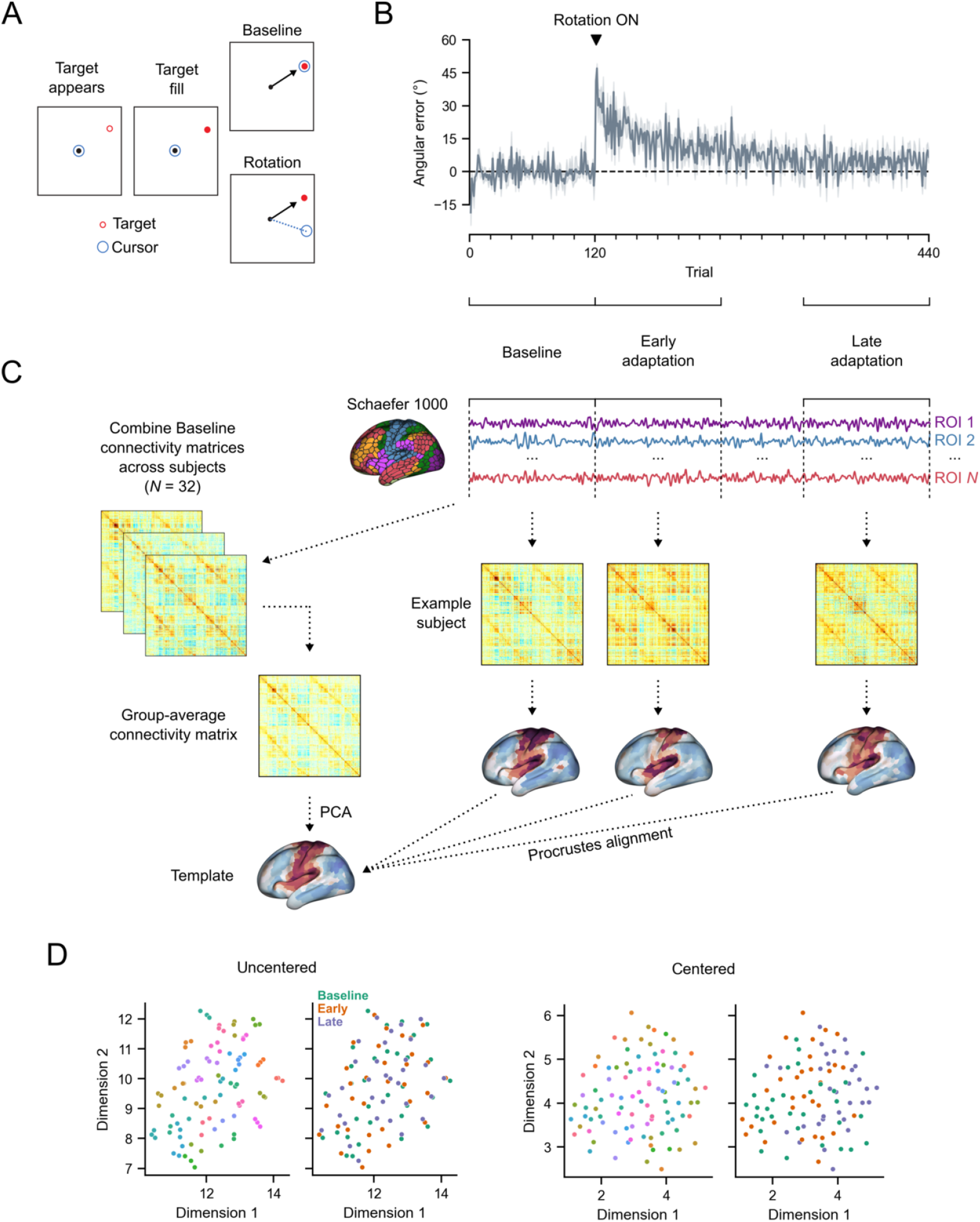
Procedure and analysis overview. (*A*) Visuomotor rotation task. On Baseline trials, cursor direction matched the aim direction. On rotation trials, cursor direction was rotated 45°clockwise relative to aim direction. (*B*) Average participant performance throughout the visuomotor rotation task. Shading indicates ±1 standard error of the mean (SEM). Three equal-length task epochs for subsequent neural analyses are indicated below: Baseline, Early adaptation (Early), and Late adaptation (Late). (*C*) Neural analysis approach. For each participant and each task epoch, functional connectivity matrices were computed using region-wise timeseries extracted with the Schaefer 1000 parcellation. Functional connectivity manifolds for each task epoch were estimated using PCA with centered and thresholded connectivity matrices (see *Materials and Methods*). All manifolds (participant ×epochs) were aligned to a common template manifold created from a group-average Baseline connectivity matrix (*left*) using Procrustes alignment. (*D*) Visualization of the similarity of connectivity matrices, both before and after centering, using UMAP. Note that uncentered connectivity matrices show strong participant-level clustering (outer-left, coloured by participant), which masks differences in task structure (inner-left, coloured by task). By contrast, centering removes this participant-level clustering (inner-right) and decouples task structure (outer-right) from these individual differences.

In order to study adaptation-related changes in functional cortical organization, we used three distinct, equal-length epochs over the time course of the task. Specifically, in addition to task Baseline (120 trials), we defined Early and Late adaptation epochs using the initial and final 120 trials, respectively, after rotation onset. For each participant, we extracted mean blood oxygenation level-dependent (BOLD) timeseries for each cortical region defined by the Schaefer 1000 parcelation [54] and estimated separate functional connectivity matrices for each epoch (Baseline, Early and Late) using the covariance matrix of the timeseries (Fig. 1C). To reduce the influence of large individual differences in functional connectivity that can obscure any task-related changes (Fig. 1D; see also [55]), all connectivity matrices were centered according to a Riemmanian manifold approach (see *Materials and Methods*; [56–58]). To demonstrate the effect of this centering procedure— and its importance for elucidating learning-related effects in the data—we projected participants’ individual covariance matrices, both before and after centring, using uniform manifold approximation (UMAP; [59]). As can be seen in Fig. 1D (*left*), prior to the centering procedure, functional network structure is dominated by participant-level clustering, which masks any task-related structure (i.e., differentiation of Baseline, Early and Late learning). However, after centering (Fig. 1D, *right*), a task structure becomes more readily apparent.

To examine reconfigurations of cortical connectivity during visuomotor adaptation, we took the centered matrices and estimated separate cortical connectivity manifolds for each participant’s Baseline, Early, and Late connectivity matrices. Using established procedures [38, 60, 61], each matrix was first transformed into an affinity matrix by computing the pairwise cosine similarity between regions after row-wise thresholding (see *Materials and Methods*). Then, we applied principal components analysis (PCA) to obtain a set of principal components (PCs), i.e. manifold, that provides a low-dimensional representation of cortical functional organization. Each matrix was then aligned to a template Baseline manifold, which was constructed using the mean of all Baseline connectivity matrices across participants (Fig. 1C, *right*). Crucially, not only did this template Baseline manifold provide a common target for manifold alignment [60], but it also allowed us to examine changes in cortical connectivity that selectively arise during the learning phase itself (i.e., Early and Late adaptation), thus increasing our sensitivity to detect deviations from the Baseline functional architecture.

### Connectivity manifold during task Baseline

The top three principal components (PCs) of the template Baseline manifold (Fig. 2A) provide a compact representation of the cortical functional organization during Baseline trials. PC 1 distinguishes somatomotor regions (positive loadings in red) from remaining cortical areas (negative loadings in blue), most prominently higher-order association regions within the default mode network (DMN), such as posteromedial cortex (PMC), as well visual areas. Meanwhile, PC 2 illustrates a gradient between visual areas and the DMN, and PC3 is a joint gradient of (i) superior-versus-inferior frontoparietal regions and (ii) lateral-versus-medial occipital regions. These top three PCs collectively explain 49.30% of the total variance (Fig. 2B). Although only the top three PCs were retained for all subsequent analyses, we note that including PC 4, which explains nearly as much variance (8.98%) as PC 3 (9.63%), does not meaningfully alter our results and interpretations (see Supplementary Fig. S1).

**Fig. 2.**
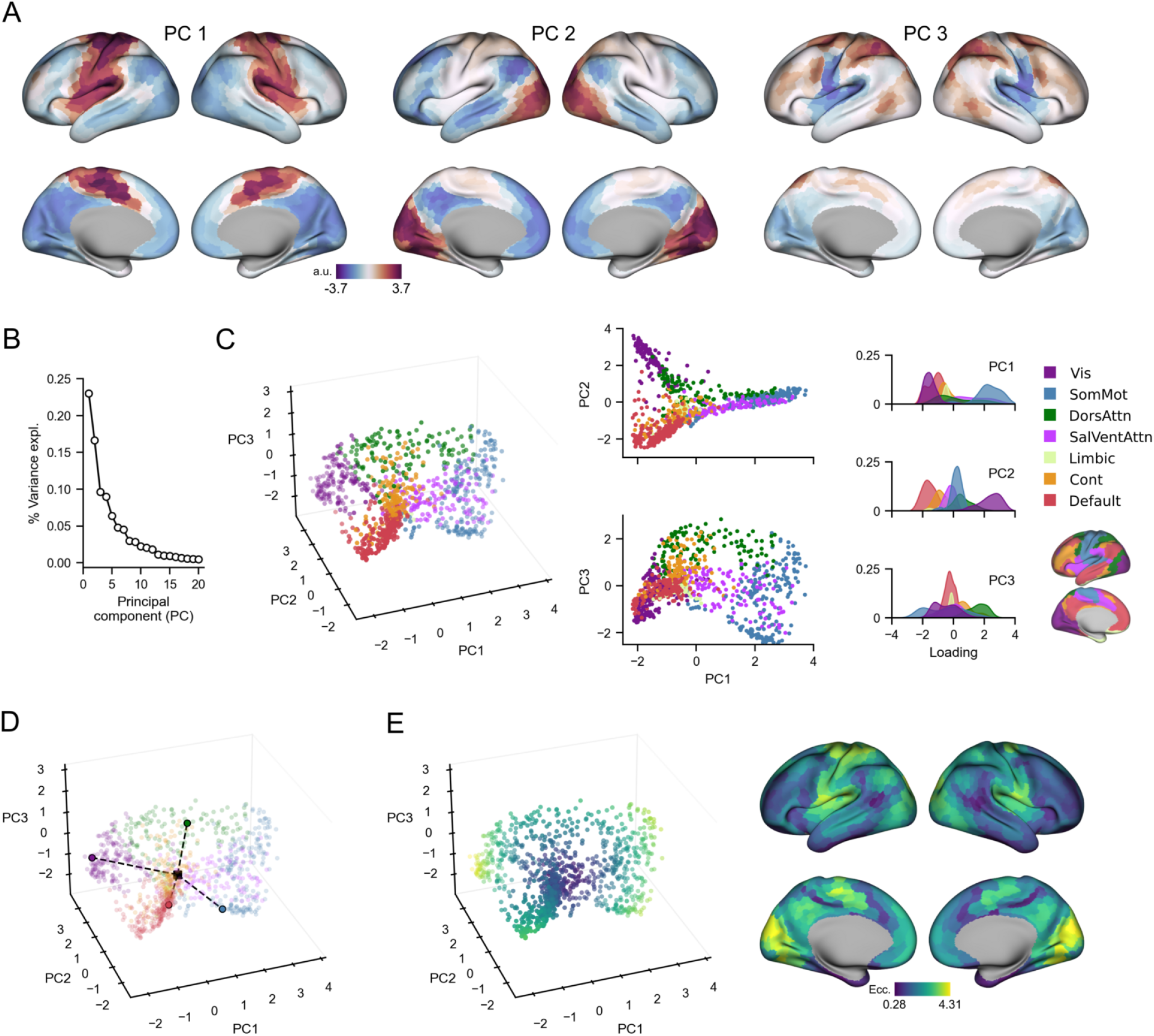
Template Baseline manifold structure and eccentricity. (*A*) Region loadings for top three PCs. (*B*) Percent variance explained for the first 20 PCs. (*C*) Functional network organization of the template Baseline manifold. *Right*, scatter plots show the embedding of each region along the top three PCs, coloured according to their intrinsic functional network [54, 62]. *Left*, probability density histograms show distribution of each functional network along each PC. Vis: Visual. SomMot: Somatomotor. DorsAttn: Dorsal attention. SalVentAttn: Salience/Ventral attention. Cont: Control. (*D*) Eccentricity calculation. Region eccentricity along the manifold is computed as the Euclidean distance (dashed line) from manifold centroid (black square). The eccentricity of four example brain regions is highlighted (bordered coloured circles). (*E*) Regional eccentricity during Baseline. Each brain region’s eccentricity is color-coded and visualized in low-dimensional space (*left*) and on the cortical surface (*right*).

Mapping brain regions onto their assigned intrinsic functional network [54, 62] shows that PCs 1 and 2 jointly differentiate visual, DMN, and somatomotor regions, resembling the tripartite structure of resting-state connectivity gradients (Fig. 2C; [38]). This differentiation is thought to reflect a fundamental feature of functional brain organization, in which the transition from largely unimodal cortex (i.e. visual and somato-motor networks) to transmodal cortex (i.e. DMN) represents a global processing hierarchy of increasing integration and abstraction from lower- to higher-order systems. [25, 38, 63]. In contrast, PC3 appears to be task-specific in that it isolates key dorsal attention, control, and somatomotor regions known to be involved in the planning and execution of hand movements required to successfully perform goal-directed actions, such as dorsal premotor cortex (PMd), superior parietal cortex (SPC), and dorsolateral prefrontal cortex (DLPFC; [64–66]).

Next, in order to characterize the relative positions of cortical brain regions along the Baseline connectivity-derived manifold space, which provides the basis for examining resultant changes in these positions throughout learning, we computed each region’s manifold eccentricity by taking its Euclidean distance from the manifold centroid (Fig. 2D; [50, 51, 67]). Eccentricity provides a multivariate index of each region’s three-dimensional embedding, in which distal regions situated at the anchors of the manifold have greater eccentricity than proximal regions within the manifold core (Fig. 2E, *left*). Highly eccentric regions therefore can be interpreted as functionally segregated from other networks in the rest of the brain, as revealed by correlating eccentricity with graph theoretical measures of integration and segregation. We find that Baseline eccentricity is positively related to node strength (*r* = 0.83, two-tailed *p <* 0.001) and within-manifold degree *z*-score (*r* = 0.49, two-tailed *<* 0.001), consistent with the idea that eccentric regions are tightly interconnected with other members of the same functional network (see Supplementary Fig. S2). Commensurate with this, we also find that eccentricity is inversely proportional to a region’s degree of cross-network integration, as measured through participation coefficient (*r* = − 0.69, two-tailed *p <* 0.001). Thus, taken together, adaptation-induced changes in a region’s functional segregation or integration can be assessed through changes in eccentricity during Early and Late adaptation.

### Manifold reconfigurations during adaptation

We found that the Early and Late adaptation epochs each exhibited distinct patterns of increases (i.e. expansion) and decreases (i.e. contraction) in manifold eccentricity relative to Baseline (Fig. 3A; for raw eccentricity maps, see Supplementary Fig. S3). To determine which regions showed significant changes in manifold eccentricity across the three task epochs (Baseline, Early and Late adaptation), we performed region-wise repeated measures ANOVAs and corrected for multiple comparisons using false-discovery rate correction (FDR; *q <* 0.05). Across cortex, we found that 131 regions showed a significant main effect of task epoch, i.e. adaptation-related changes, with 111 of these regions forming 14 contiguous clusters (Fig. 3B). Major clusters include contiguous regions spanning from left (contralateral) PMd to SPC (18 regions), left PMC (20 regions), and dorsolateral portions of bilateral extrastriate cortex (left = 22 regions; right = 11 regions). Smaller clusters and singleton regions were also observed throughout the rest of cortex, and the combination of all clusters/regions spanned all six non-Limbic functional networks (Fig. 3C). Note that these topographical clusters arise because of the large degree of spatial autocorrelation along each dimension (see Fig. 2A). That is, topographically adjacent regions are more likely to have similar connectivity profiles, and thus have similar projections onto the manifold.

**Fig. 3.**
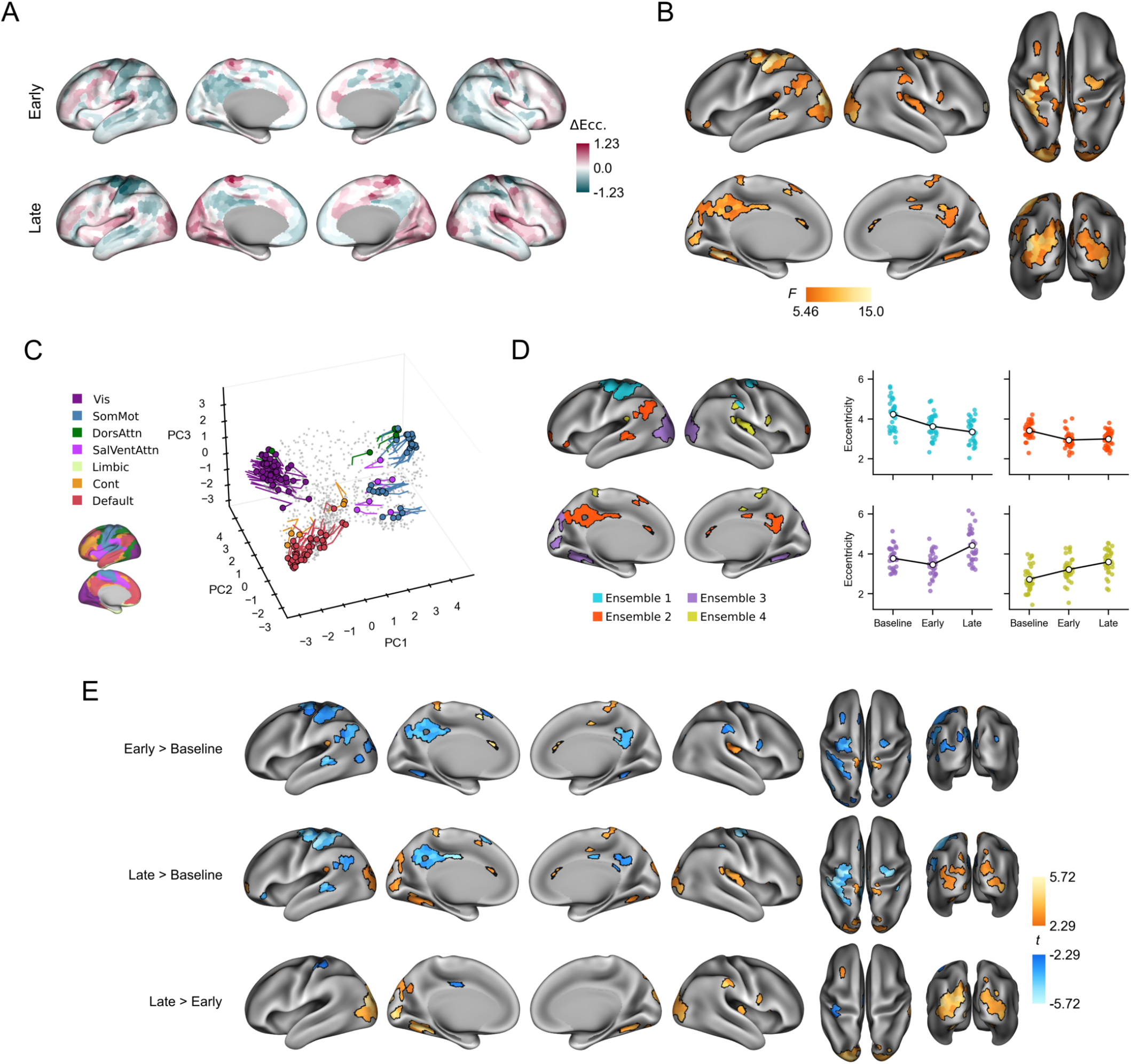
Adaptation-related changes in manifold eccentricity. (*A*) Region-wise mean changes in eccentricity during Early and Late adaptation, relative to Baseline. Positive (mauve) and negative (teal) values indicate relative increases and decreases in eccentricity, respectively. (*B*) Significant changes in eccentricity across task epochs according to region-wise repeated measures ANOVAs with false-discovery rate (FDR) correction for multiple comparisons (*q <* 0.05). (*C*) Temporal trajectories of statistically significant regions from *B* in low-dimensional space. Coloured circles indicate each region’s initial position during Baseline and the traces show the unfolding displacement of that region during Early and Late adaptation. Each region is coloured according to its functional network assignment (*left*). Nonsignificant regions are shown in gray point cloud. (*D*) Patterns of effects for four ensembles of significant regions in *B* derived from *k*-means clustering on each brain region’s coordinates during Baseline (see *C*). Scatter plots (*right*) show within-ensemble mean eccentricity for each participant, and line plot overlays (white markers) show the group mean across task epochs. (*E*) Pairwise contrasts of eccentricity between task epochs. Region-wise paired *t*-tests were performed for each contrast and FDR correction was applied across all comparisons (*q <* 0.05). Positive (orange) and negative (blue) values show significant increases and decreases in eccentricity, respectively.

To provide a concise summary of the ANOVA results presented above, we used *k*-means clustering to group regions with significant main effects according to their coordinates at Baseline (Fig. 3C, *coloured circles*). This approach gave way to brain regions that tended to exhibit similar temporal trajectories in manifold space during adaptation (Fig. 3C *traces*). The clustering analysis revealed four ensembles of regions (Fig. 3D): Ensemble 1 (blue) is composed of somatomotor and dorsal attention network regions that load positively onto PC 1 and 3, which includes left sensorimotor regions that make up the largest cluster in Fig. 3B, along with right PMd and parietal regions; Ensemble 2 (red) primarily involves higher-order transmodal areas of the DMN that load negatively onto PC 1 and 2, such as PMC, angular gyrus (AG), and superior temporal sulcus (STS); Ensemble 3 (purple) is mainly comprised of visual regions, which include bilateral extrastriate and parahippocampal regions, which load negatively and positively onto PC 1 and 2, respectively; and Ensemble 4 (yellow), which includes remaining regions in somatomotor and salience/ventral attention networks that load positively on PC1 but negatively on PC3. Computing the average eccentricity of each ensemble reveals distinct patterns of contractions and expansions along the manifold that characterize the key changes in connectivity during adaptation (Fig. 3D, *right*).

Next, we directly examined the region-based changes in eccentricity between each task epoch by performing follow-up paired *t*-tests on the regions that exhibited significant main effects (in Fig. 3B), with corrections for multiple comparisons applied across all tests using FDR correction (*q <* 0.05; Fig. 3E). As revealed by a contrast of Early*>*Baseline, Early adaptation is primarily characterized by reductions in eccentricity, i.e. manifold contractions, of regions belonging to Ensembles 1 and 2, including regions in somatomotor and premotor cortex, as well as areas of the DMN, such as bilateral PMC and angular gyrus (AG). Although regions in Ensemble 3 (visual network) collectively trend towards manifold contraction during Early learning (see Fig. 3D), only eight regions in extrastriate and parahippocampal cortices exhibited significant contractions, after corrections for multiple comparisons. Meanwhile, regions in Ensemble 4 showed significant increases in eccentricity over the same time window, i.e. manifold expansion.

As revealed by the contrast of Late*>*Baseline (Fig. 3E, *middle*), we found that sensorimotor and DMN regions in Ensembles 1 and 2, respectively, maintained their contraction during Late adaptation. Performing a Late*>*Early contrast (Fig. 3E, *bottom*) shows that the extent of the contraction in these regions did not significantly differ between epochs, with the exception of an increased contraction in left somatosensory cortex and a subregion within left PMC. However, by and large, the main characteristic feature of Late adaptation is the expansion of visual cortex along the manifold, including bilateral extrastriatal regions. These effects are more pronounced for the Late*>*Early contrast than for the Late*>*Baseline contrast, which is a result of the overall trend towards manifold contraction of these visual areas areas during Early adaptation.

Taken together, the above pattern of results suggest that, during Early adaptation, several visual, sensorimotor and transmodal areas in the DMN begin to integrate with regions outside of their respective functional networks. By contrast, during Late adaptation when performance plateaus and errors become minimized, our findings suggest that visual cortical regions, particularly higher order visual areas, become functionally segregated from other brain networks. In the next section, we seek to directly test these interpretations of manifold contractions and expansions during Early and Late adaptation, respectively.

### Connectivity changes underlying manifold reconfigurations

Because eccentricity represents a multivariate measure of a region’s overall connectivity profile, we performed seed connectivity analyses in order to help characterize the changes in connectivity that underlie the patterns of manifold contraction and expansion we observed throughout adaptation. To describe connectivity changes during Early adaptation, we selected representative regions of the three largest clusters in the Early*>*Baseline contrast, which included left PMC, left SPC, and left PMd (see *Materials and Methods*). For each region, we contrasted seed connectivity maps between the Early and Baseline epochs (Early*>*Baseline) by computing regionwise paired *t*-tests, producing contrast maps for each seed region (Fig. 4A). We show the unthresholded contrast maps to allow visualization of the complete array of connectivity differences that collectively contribute to the eccentricity changes.

**Fig. 4.**
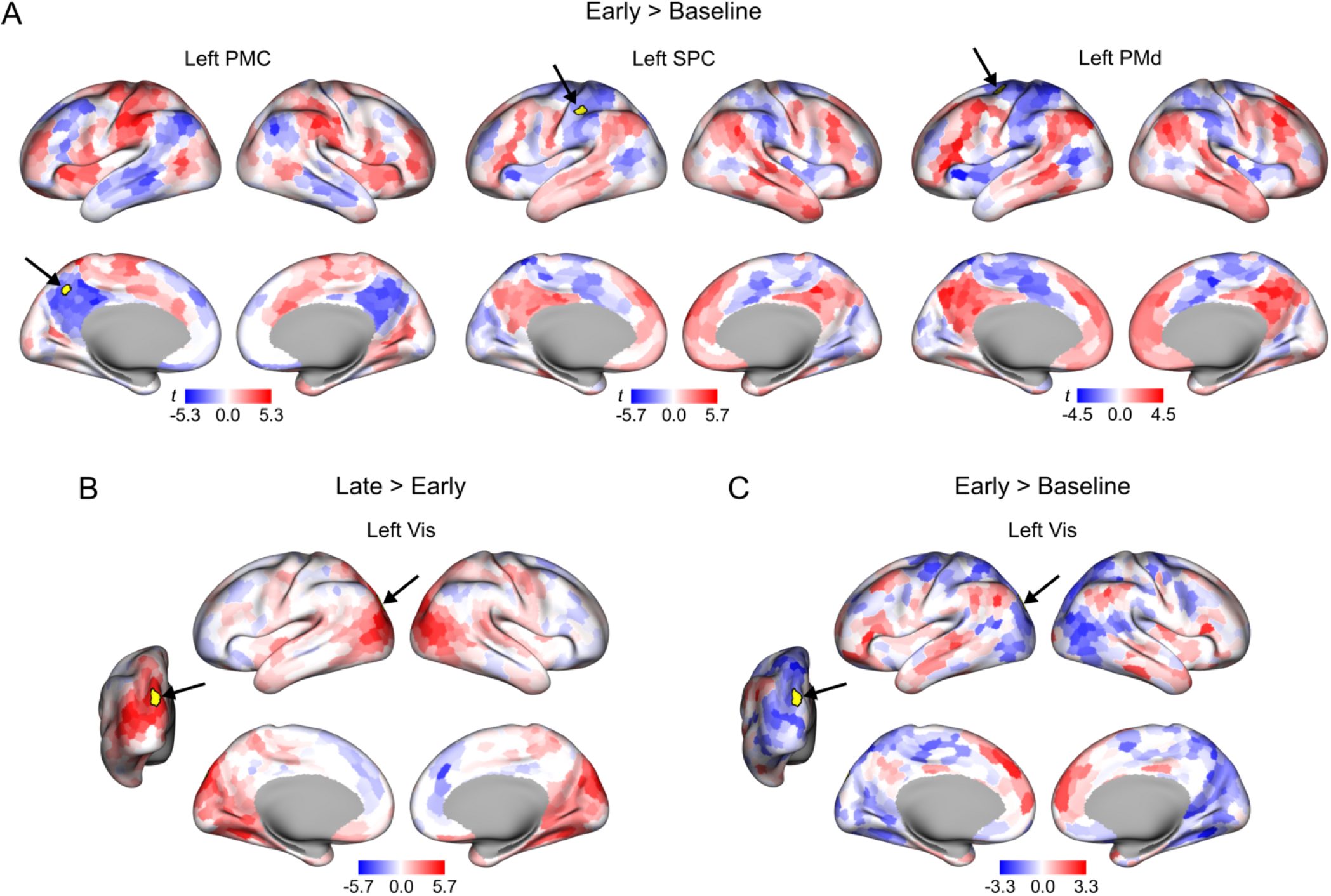
Patterns of connectivity differences underlying changes in manifold eccentricity. (*A*) Early*>*Baseline seed connectivity contrast maps for left PMC, SPC, and PMd. Selected seed regions are shown in yellow and are also indicated by arrows. Positive (red) and negative (blue) values show increases and decreases in connectivity, respectively, during Early adaptation relative to Baseline. (*B*) Late*>*Early and (*C*) Early*>*Baseline seed connectivity contrast maps for the left visual/extrastriate (Vis) seed region.

During Early adaptation, we found that the left PMC seed region exhibited decreased connectivity with other PMC subregions across both hemispheres, as well as with other DMN areas located in bilateral AG and STS, and with left dorsolateral prefrontal cortex (DLPFC). Instead, the PMC exhibited increased connectivity with sensorimotor regions such as PMd and SPC, along with anterior portions of frontal cortex and insula. Notably, the opposite pattern can be observed for both the left SPC and PMd seed regions, which exhibited decreased connectivity with other sensorimotor regions in favour of increasing their connectivity with areas of the DMN (e.g., bilateral PMC, AG, STS) and the DLPFC. Together, these findings indicate that manifold contractions of the PMC, SPC, and PMd during Early adaptation largely arise from increased integration between sensorimotor regions (Ensemble 1) and higher-order transmodal regions of the DMN (Ensemble 2).

We also repeated the seed connectivity analysis to investigate the basis of the manifold expansion of visual cortex observed during Late adaptation. Using the Late*>*Early eccentricity contrast, we selected a representative region from the left extrastriate cluster, which was the largest cluster across both hemispheres. By contrasting seed maps between the Late and Early epochs (i.e. Late*>*Early), we found that connectivity increased within bilateral visual cortex and parahippocampal regions (Fig. 4B). Meanwhile, connectivity to the rest of cortex remained relatively unchanged, with the exception of subtle connectivity reductions in dorsomedial frontal cortex. These results suggest that the segregation/expansion of visual areas during Late adaptation is mainly driven by increased intraconnectivity of visual cortex rather than a decoupling from the rest of cortex. Notably, this same visual seed region also exhibited a significant contraction in our Early*>*Baseline eccentricity contrast (see Fig. 3E), and as such, we additionally contrasted this region’s seed connectivity during the Early and Baseline epochs. This analysis revealed decreased connectivity with visual and sensorimotor regions during Early adaptation, while also showing increased connectivity with DMN areas, such as AG, STS, and dorsomedial frontal cortex (Fig. 4C). Thus, consistent with the pattern of effects shown above for areas PMC, SPC and PMd, we found that greater connectivity with the DMN also underlies significant manifold contractions of visual areas. Together, this suggests that increased functional interactions between unimodal and transmodal cortical areas is a feature of early learning.

### Eccentricity relates to performance during Early adaptation

Thus far, we have characterized within-participant alterations to manifold structure throughout adaptation, revealing patterns of manifold contraction and expansion expressed across individuals. It is well-established, however, that significant intersubject variability exists during the initial phases of learning, when visual-motor errors are largest [17–19]. Consistent with this prior work, we find a large degree of between-participant variability in performance during Early adaptation (Fig. 5A), which we measured by computing the median angular error for each participant (i.e. Early error; Fig. 5B). Given these prominent individual differences in performance, we next asked whether this inter-subject variability is related to manifold structure during Early adaptation, as captured by eccentricity.

**Fig. 5.**
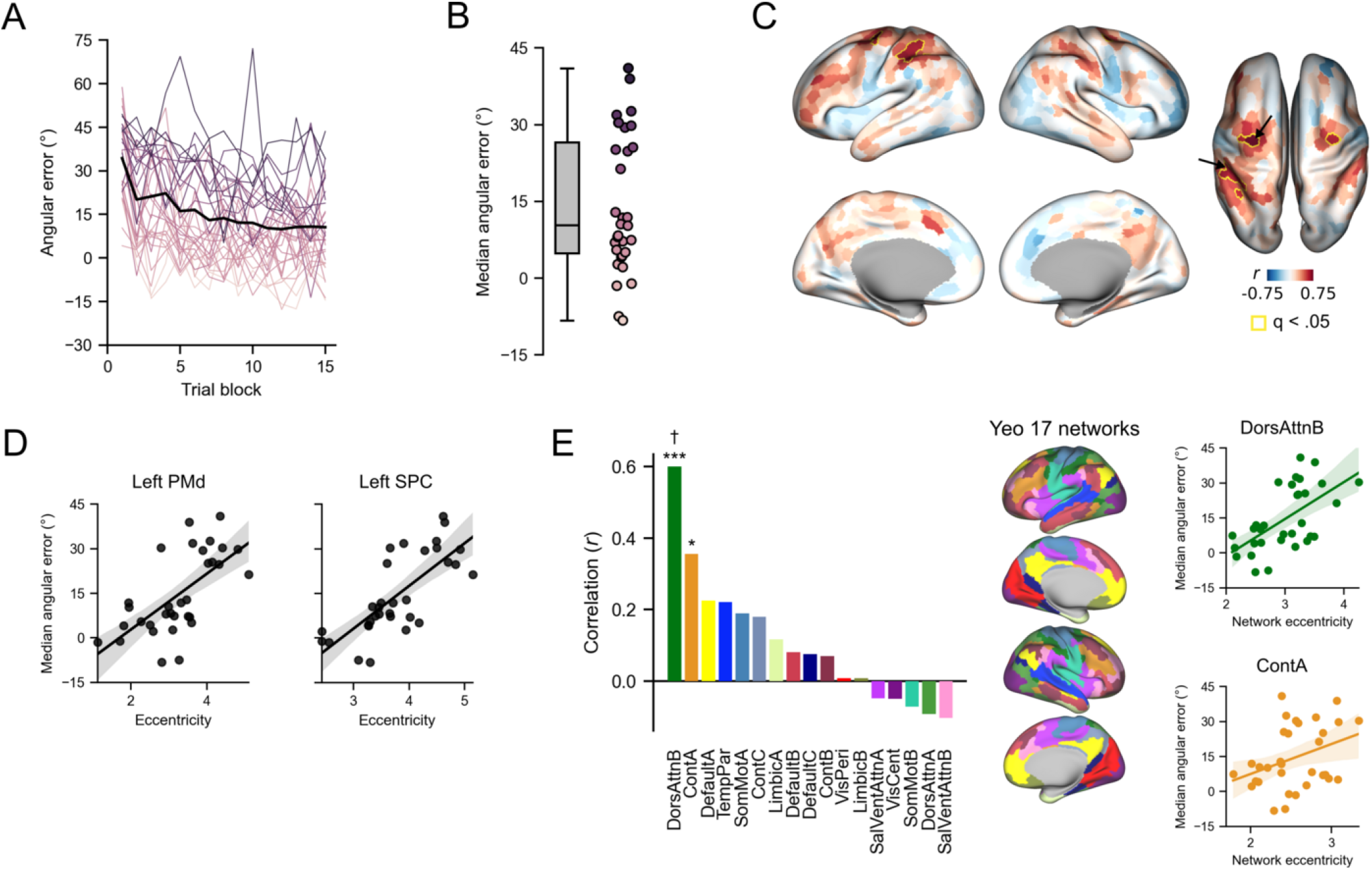
Manifold eccentricity during Early adaptation relates to task performance. (*A*) Individual differences in behavioural performance during Early adaptation. Black line shows the average error across participants, binned by trial block, and coloured traces show binned error for individual participants. Different participants are coloured according to their median error in *B*. (*B*) Distribution of median error during Early adaptation. Colour in scatter plot indicates median error, with lighter colour in scatter plot indicating lower Early error (better learning). (*C*) Correlation map between Early eccentricity and Early error. Yellow traces show significant regions following FDR correction (*q <* 0.05). Arrows indicate left PMd and SPC seed regions in Fig. 4A, which are used as exemplar significant regions in *D*. (*D*) Example correlations from left PMd and SPC seed regions in Fig. 4A. (*E*) Correlations between network eccentricity and Early error (*left*); scatter plots are shown for the two largest correlations (*right*). Bar and scatter plot colours correspond to network colours shown on brain surface (*middle*). Of all networks, Dorsal Attention B and Control A showed the strongest correlations (*right*). VisCent: Visual Central. VisPer: Visual Peripheral. SomMotA: Somatomotor A. SomMotB: Somatomotor B. TempPar: Temporal Parietal. DorsAttnA: Dorsal Attention A. DorsAttnB: Dorsal Attention B. SalVentAttnA: Salience/Ventral Attention A. SalVentAttnB: Salience/Ventral Attention B. ContA: Control A. ContB: Control B. ContC: Control C. \**p <* 0.05, \*\*\**p <* 0.001, *†q <* 0.05

To examine this question at the region-level, we calculated the correlation between participants’ Early error and the eccentricity values within each cortical region during Early adaptation (Fig. 5C). Following FDR-correction for multiple comparisons (*q <* 0.05), we found that regions within left (contralateral) parietal cortex and bilateral PMd exhibited significant positive associations between their manifold eccentricity and participant Early error (i.e. greater eccentricity corresponds with greater error or worse performance). Note that, across participants, these same regions exhibit overall manifold contractions during Early adaptation (see Fig. 3E), and thus participants with greater contractions (i.e. lower eccentricity) in these regions during Early adaptation show faster learning (i.e. lower Early error). Also recall that manifold contractions of these same regions (the left PMd and SPC seed regions used for connectivity analyses (Fig. 5C, *arrows*) are associated with increased connectivity with higher-order transmodal regions within the DMN (Fig. 4A). Taken together, these findings suggest that participants who adapt more rapidly express a greater degree of integration between sensorimotor and higher-order association networks during Early adaptation. This is consistent with the idea that the coupling of transmodal and sensorimotor cortical regions during adaptation reflects the recruitment of explicit learning processes, which exert top-down control over the sensorimotor system [16–18].

It is noteworthy that the region-level correlations shown in Fig. 5C exhibit a high degree of spatial contiguity; that is, the parietal and premotor cortical regions that pass FDR-correction (noted above) are situated within much larger clusters of regions that exhibit a very similar pattern of correlations with learning performance. This topography suggests that the association between manifold eccentricity and participant behaviour during Early adaptation may be further characterized at the level of distributed functional networks. To explore this possibility, we mapped each region onto its respective functional network and, within each participant, computed the average manifold eccentricity for each network (i.e. network eccentricity). For this purpose, we used the 17-network mapping in order to capitalize on the improved spatial precision—and thus ability to better localize effects—compared to the 7-network mapping [54, 62]. Next, we correlated, for each network, its eccentricity during Early adaptation with participants’ Early error. Among these networks, the Dorsal

Attention B (*r* = 0.60, two-tailed *p <* .001; Fig. 5E) and Control A (*r* = 0.35, two-tailed *p* = 0.046) networks showed a significant positive association with Early error (although the Control A did not survive FDR-correction, *<* 0.05). Collectively, these two networks span several parietal (e.g., SPC, intraparietal sulcus), premotor (e.g., PMd, frontal eye fields), and prefrontal areas (e.g., DLPFC), which together represent an array of brain areas previously implicated in higher-order sensorimotor processing and the top-down control of goal-directed behaviour [65, 68–70].

## Discussion

Complex behaviour depends on the coordinated operation of several specialized neural systems distributed throughout the brain. During sensorimotor adaptation, these distributed systems must modify their interactions to ensure that motor behaviour appropriately responds to changes in environmental dynamics and regularities. While much focus to date has been on understanding the cerebellar-dependent mechanisms that underlie sensorimotor adaptation, our understanding of the contributions and functional reorganization of cortical systems remains incomplete [71]. Here, we capitalized on recent analytical methods that link together topographic and functional brain organization [38, 60] in order to quantify adaptation-related changes in cortical activity patterns, and how features of this reorganization relate to learning performance.

By projecting subjects’ cortical functional connectivity patterns into compact low-dimensional manifold spaces, we found that adaptation was primarily characterized by increasing manifold contractions of higher-order sensorimotor regions in parietal and premotor cortex, as well as transmodal areas of the DMN. Further analyses revealed that these manifold contractions were the result of greater covariance in neural activity between transmodal (i.e., DMN and frontoparietal networks) and unimodal (i.e. sensorimotor) systems, which was largely maintained across the entire adaptation period. In addition, we found that, by the late stages of adaptation, when visual-motor errors had been largely reduced, visual cortical regions exhibited expansion along the cortical manifold, a pattern that was explained by greater intraconnectivity within visual cortex. Finally, our analyses revealed that these changes have important behavioural correlates, as faster overall adaptation was linked to increased covariance between sensorimotor and transmodal areas of the DMN. Together, our results provide a novel characterization of the macroscale cortical changes that support human sensorimotor adaptation and performance. As we discuss below, these findings have important implications for our understanding of the cortical basis of adaptation, in general, and the role that association cortex, and the DMN in particular, plays in the organization of adaptive behaviour.

Several prior fMRI studies have revealed adaptation-related increases and decreases in the activity of individual sensorimotor cortical brain regions, including areas in motor, premotor, and parietal cortex [3–5, 7, 23, 72]. Our results expand on these findings by suggesting that these individual region-based changes are part of a broader reorganization of the cortical landscape that occurs during adaptation. Specifically, our analyses suggest that sensorimotor areas become increasingly integrated with higher-order association areas in the DMN and DLPFC following a visuomotor perturbation. Contemporary models of cortical organization [25, 38] suggest that transmodal regions are important for organizing behaviour in an increasingly abstract manner. It is possible that the observed changes in the cortical landscape, therefore, reflect the increased need for more abstract control over unimodal systems. This perspective is consistent with behavioural and lesion work indicating that adaptation recruits cortically-driven explicit learning processes (e.g., mental rotation, working memory, etc.) that are strategic and declarative in nature, and thus presumed to involve brain areas in higher-order association cortex (e.g., prefrontal regions; [15, 24]). Furthermore, existing evidence suggests that faster learning across participants results from the greater recruitment of explicit learning processes during adaptation [19]. Consistent with this, we find that faster learning across participants is associated with a greater manifold contraction of higher-order sensorimotor regions in parietal and premotor cortex, and that these contractions reflect the increased covariance of these areas with regions of the DMN and prefrontal cortex (Fig. 4). We find it noteworthy that these parietal and premotor cortical areas belong to the dorsal attention (DAN) network (see Fig. 5E), given that this network, in particular, has been heavily implicated in the top-down control of attention and action [62, 68, 70]. However, we recognize that, although our study highlights the interactions of both unimodal and transmodal systems during learning, our design does not allow us to delineate the specific higher-order control processes that this pattern reflects.

Our data also have important implications for understanding the role of the DMN in cognition and behaviour. Traditionally, the DMN was thought to mainly support perceptually decoupled states, such as mind-wandering, mental time travel or autobiographical memory [35, 73, 74]. However, recent studies have shown that regions in this system can also play a direct role in task-based cognition, particularly under situations where decision-making cannot be based on immediate sensory input and must instead be based on prior information (e.g., from a previous trial) [28–31, 36, 75]. Our study adds to this emerging literature by providing the first evidence that the DMN is also involved in a sensorimotor process like error-based learning, whereby action selection on the current trial (i.e., what direction to move) must be based on information accrued across previous trials (e.g., a memory of visual-motor errors experienced [76]. As such, our findings are inconsistent with the view that the DMN is strictly ‘task-negative’, but rather that its activity may be important for organizing cognition and behaviour in a more abstract, flexible manner [35]. This process can be important not only when we are engaged in stimulus-independent patterns of thought, such as when we daydream or imagine the future, but also in more coupled modes of cognition, such as when adapting one’s current motor behaviour based on the history of sensory feedback experienced [76].

One curious observation was our finding that the visual cortex exhibited contraction along the cortical manifold during Early adaptation whereas, during Late adaptation, it exhibited expansion (Fig. 3E). Our further analyses indicated that this reversal pattern resulted from the relative increase in covariance, during Early adaptation, between visual cortex and areas of the DMN (e.g., medial prefrontal cortex, angular gyrus, superior temporal gyrus) compared to a relative decrease in this covariance during Late adaptation. While we can only speculate on the nature of these changes, one possibility is that they reflect a relative shift in the neural processing of visual errors experienced by participants across the different phases of learning. During Early adaptation, subjects experience large visual errors (45°), which tend to engage explicit re-aiming processes to help minimize those errors [77]. At the neural level, this would require that errors sensed by the visual system be fed forward to higher-order association cortex, which, in turn, would implement a re-aiming strategy to help reduce those visual errors. This would presumably be manifest as increased covariance between visual and transmodal cortex, which is consistent with our Early adaptation results. Likewise, by the end of learning, when visual errors have been reduced to near baseline levels, the feed forward exchange of information from visual to transmodal cortex would be expectantly reduced. This would presumably be manifest as decreased covariance between visual and transmodal cortex, which is also consistent with our late adaptation results.

Another, albeit not mutually exclusive possibility, is that the pattern of manifold expansion-then-contraction of visual cortex described above relates to learning-dependent changes in the top-down modulation of visual cortical activity by transmodal cortex. For instance, during Early adaptation, when visual errors are large and numerous (and when learning rates are maximal), the attentional processing of visual errors is likely to be heightened and prioritized as compared to during Late adaptation, when errors tend to be much smaller in magnitude and when performance has more or less stabilized. Prior work has shown that the allocation of attentional resources during learning plays a critical role in successful sensorimotor adaptation [78–80] and, similarly, that the allocation of spatial attention during motor planning modulates neural activity in visual cortex [81–83]. Although the neural circuits by which visual cortex is modulated during tasks involving motor learning and control remain poorly understood, such modulation likely involves top-down projections from higher-order brain areas in association cortex [84, 85]. Taken together, our visual cortex findings are likely to be explained, at least in part by, both bottom-up and top-down interactions between transmodal and visual cortex.

In summary, here we applied recent dimensionality reduction approaches in order to describe the changing functional architecture of cortex during sensorimotor learning. This approach enabled us to identify adaptation-related shifts in low-dimensional connectivity structure that are driven by increasing integration between regions within sensorimotor and higher-order association networks, and later in adaptation, functional segregation of visual areas. These findings offer a unique perspective in our understanding of the cortical contributions to sensorimotor adaptation, which not only have important implications in contemporary theories of motor learning, but also the role of transmodal cortex in task-based performance.

## Acknowledgements

This work was supported by operating grants from the Canadian Institutes of Health Research (CIHR) awarded to J.P.G (MOP126158). J.P.G. was also supported by a NSERC Discovery Grant, as well as funding from the Canadian Foundation for Innovation. D.J.G was supported by a Natural Sciences and Engineering Research Council (NSERC) graduate award. The authors thank Martin York, Sean Hickman, Don O’Brien, and Michael Lewis for technical assistance.

## Author Contributions

D.J.G.: Conceptualization, Methodology, Data Analysis, Software, Writing (original draft, visualization). C.N.A.: Methodology, Writing (review). D.P.S. Data collection, Writing (review). J.Y.N.: Data collection. R.D.M.: Methodology, Software. J.R.F.: Supervision, Conceptualization, Funding. J.P.: Methodology, Writing (original draft, review). J.P.G. Supervision, Conceptualization, Funding, Methodology, Writing (original draft, review).

Authors declare no competing interests. R.D.M. is currently employed by Octave Bioscience, which has no relation to the present research.

## Materials and Methods

### Participants

40 right-handed individuals (13 males) between the ages of 18 and 35 (*M* = 22.5, *SD* = 4.51) participated in the study. Eight participants were excluded based on in-scanner head motion (four participants; *>*2mm translation or *>*2^◦^ rotation in a single scan) or inability to correctly perform and learn the visuomotor rotation task (four participants). Right handedness was assessed using the Endinburgh Handedness Questionnaire [86]. Participants’ written, informed consent was obtained before commencement of the experimental protocol. The Queen’s University Research Ethics Board approved the study and it was conducted in accordance with the principles outlined in the Canadian Tri-Council Policy Statement on Ethical Conduct for Research Involving Humans and the principles of the Dec-

### Experimental setup

Participants performed hand movements directed towards a target by applying a directional force onto an MRI-compatible force sensor (ATI Technologies) using their right index finger and thumb. The target and cursor stimuli were rear-projected with an LCD projector (NEC LT265DLP projector, 1024 × 768 resolution, 60 Hz refresh rate) onto a screen mounted behind the participant. The stimuli on the screen were viewed through a mirror fixed to the MRI coil directly above participants’ eyes, thus preventing participants from being able to see the hand. The force sensor and associated cursor movement were sampled at 500Hz.

### Visuomotor rotation task

We used a well-established motor learning task, the visuomotor rotation task [53], to probe sensorimotor adaptation. To start each trial, the cursor (20 pixel radius) appeared in a central start position (25 pixel radius). A white target circle (30 pixel radius) appeared at one of eight locations (0^◦^, 45^◦^, 90^◦^, 135^◦^, 180^◦^, 225^◦^) on an invisible ring around the central position (300 pixel radius) and filled in (white) following a 200 ms delay. Once launched, the cursor would travel the 300 pixel distance to the ring over a 750 ms period (with a bell-shaped velocity profile) before becoming stationary at the ring to provide participants with end-point error feedback. If the cursor overlapped with the target to any extent, the target would become green, signifying a target “hit". Each trial was separated by 4 s and within this period, participants had 2.6 s from target presentation to complete the trial (including the 200 ms target delay, participants’ own reaction time, and the 750 ms cursor movement; any remaining time was allotted to providing the end-point error feedback). At 2.6 s the trial was ended, the screen was blanked, the data saved, and participants would briefly wait for the next trial to begin. Reaction times were not stressed in this experimental procedure. On trials in which the reaction time exceeded 2.6 s, the trial would end, and the participant would wait for the next trial to begin. These discarded trials were rare (0.56% across all trials, all participants) and were excluded from behavioural analyses, but were kept in the neuroimaging analysis due to the continuous nature of the fMRI task and our focus on functional connectivity analyses. On each trial we measured the angular error between the target and the final cursor position. Trials with reactions times *<*100 ms or *>*2000 ms were discarded (the former value was chosen as a conservative threshold on prepotent or anticipatory responses).

### Procedure

Participants performed the visuomotor rotation task during two identical fMRI sessions, separated by exactly 24 hours. In each session, participants completed a single continuous task scan (29 minutes and 52 seconds), which comprised of 120 baseline trials (15 sets of 8 trials) with no cursor rotation (baseline), followed by 320 rotation trials, in which a 45^◦^ clockwise rotation of the cursor was applied. In a subsequent washout scan (8 minutes and 32 seconds), conditions were restored to baseline (i.e. no rotation of cursor) and participants performed 120 washout trials. We also interspersed three 6-minute resting state scans before and after the task scan, and after the washout scan. During these resting-state scans, participants were instructed to rest with their eyes open, while fixating a central cross location presented on the screen.

Note that the aforementioned procedure, repeated over two fMRI sessions, was collected to explore individual differences in functional brain organization related to sensorimotor adaptation, de-adaptation, and subsequent re-adaptation (see [57, 87]). Given that the present study aims to specifically examine changes in functional brain architecture during initial adaptation to a novel visuomotor perturbation, we focused our analyses exclusively on the task scans of the first session.

### MRI acquisition and preprocessing

Participants were scanned using a 3T Siemens TIM MAGNETOM Trio MRI scanner located at the Centre for Neuroscience Studies, Queen’s University (Kingston, Ontario, Canada). For each participant on each day, we collected a T1-weighted ADNI MPRAGE anatomical (TR = 1760 ms, TE = 2.98 ms, field of view = 192 mm × 240 mm × 256 mm, matrix size = 192 × 240 × 256, flip angle = 9°, 1 mm isotropic voxels). Functional MRI volumes were acquired using a 32-channel head coil and a T2*-weighted single-shot gradient-echo echo-planar imaging (EPI) acquisition sequence (TR = 2000 ms, slice thickness = 4 mm, in-plane resolution = 3 mm × 3 mm, TE = 30 ms, field of view = 240 mm × 240 mm, matrix size = 80 × 80, flip angle = 90°), and acceleration factor (integrated parallel acquisition technologies, iPAT) = 2 with generalized autocalibrating partially parallel acquisitions (GRAPPA) reconstruction. Each volume comprised 34 contiguous (no gap) oblique slices acquired at a 30° caudal tilt with respect to the plane of the anterior and posterior commissure (AC-PC), providing whole-brain coverage. For the task scan, we collected a single, continuous scan of 896 imaging volumes. This included an additional 8 imaging volumes at both the beginning and the end of the experimental run.

Preprocessing of anatomical and functional MRI data was performed using *fMRIPrep* 20.1.1 ([88, 89]; RRID:SCR_016216) which is based on *Nipype* 1.5.0 ([90, 91]; RRID:SCR_002502). Many internal operations of *fMRIPrep* use *Nilearn* 0.6.2 [92, RRID:SCR_001362], mostly within the functional processing workflow. For more details of the pipeline, see the section corresponding to workflows in *fMRIPrep*’s documentation. Below we provide a condensed description of the preprocessing steps.

T1w images were corrected for intensity non-uniformity (INU) with N4BiasFieldCorrection [93], distributed with ANTs 2.2.0 [94, RRID:SCR_004757]. The T1w-reference was then skull-stripped with a *Nipype* implementation of the antsBrainExtraction.sh workflow (from ANTs), using OASIS30ANTs as target template. Brain tissue segmentation of cerebrospinal fluid (CSF), white-matter (WM) and gray-matter (GM) was performed on the brain-extracted T1w using fast [FSL 5.0.9, RRID:SCR_002823, 95].A T1w-reference map was computed after registration of 2 T1w images (after INU-correction) using mri_robust_template [FreeSurfer 6.0.1, 96]. Brain surfaces were reconstructed using recon-all [FreeSurfer 6.0.1, RRID:SCR_001847, 97], and the brain mask estimated previously was refined with a custom variation of the method to reconcile ANTs-derived and FreeSurfer-derived segmentations of the cortical gray-matter of Mindboggle [RRID:SCR_002438, 98]. Volume-based spatial normalization to standard space (MNI152NLin6Asym) was performed through non-linear registration with antsRegistration (ANTs 2.2.0), using brain-extracted versions of both T1w reference and the T1w template.

For each BOLD run, the following preprocessing was performed. First, a reference volume and its skull-stripped version were generated using a custom methodology of *fMRIPrep*. Head-motion parameters with respect to the BOLD reference (transformation matrices, and six corresponding rotation and translation parameters) are estimated before any spatiotemporal filtering using mcflirt [FSL 5.0.9, 99]. BOLD runs were slice-time corrected using 3dTshift from AFNI 20160207 [100, RRID:SCR_005927]. The BOLD reference was then co-registered to the T1w reference using bbregister (FreeSurfer) which implements boundary-based registration [101]. Co-registration was configured with six degrees of freedom. The BOLD timeseries were resampled with *a single interpolation step* by composing all the pertinent transformations (i.e. head-motion transform matrices, and co-registrations to anatomical and output spaces). BOLD timeseries were resampled onto their original, native space, as well as standard space (MNI152NLin6Asym), using antsApplyTransforms (ANTs), configured with Lanczos interpolation to minimize the smoothing effects of other kernels [102]. Subcortical and cerebellar data in standard space was combined with resampled BOLD timeseries on the *fsaverage* surface to produce *Grayordinates* files [103] containing 91k samples, using fsaverage as the intermediate standardized surface space. Resampling onto *fsaverage* was performed using mri_vol2surf (FreeSurfer).

A set of 34 motion and physiological regressors were extracted in order to mitigate the impact of head motion and physiological noise. The six head-motion estimates calculated in the correction step were expanded to include temporal derivatives and quadratic terms of each of the original and derivative regressors, totalling 24 head-motion parameters [104]. 10 component-based physiological regressors were estimated using the aCompCor approach [105, 106], where the top five principal components were separately extracted from WM and CSF masks. Principal components were estimated after high-pass filtering the preprocessed BOLD timeseries (using a discrete cosine filter with 128s cut-off).

### Region timeseries extraction

The first three imaging volumes were discarded to avoid scanner equilibrium effects. Then, for each participant and scan, the average BOLD timeseries were computed from the grayordinate timeseries for each of the 998 regions defined according the Schaefer 1000 parcellation ([54]; two regions are removed from the parcellation due to their small parcel size). The Schaefer 1000 parcellation was selected in order to balance computational feasibility across participants and task epochs (see below) with high spatial resolution. Region timeseries were denoised using the above-mentioned confound regressors in conjunction with the discrete cosine regressors (128s cut-off for high-pass filtering) produced from *fMRIprep* and low-pass filtering using a Butterworth filter (100s cut-off) implemented in *Nilearn*. Finally, all region timeseries were *z*-scored.

### Functional connectivity estimation

For every participant, region timeseries from the task scan were spliced according to four equal-length task epochs (each 240 imaging volumes). Baseline comprised of the initial 120 trials with the veridical hand-to-cursor motion mapping (i.e. no cursor rotation), whereas Early and Late adaptation consisted of the first and last 120 trials after rotation onset, respectively. Washout was defined as the 120 trials without cursor rotation in the washout scan. Then, we separately generated functional connectivity matrices for each epoch by computing the region-wise covariance matrix using the Ledoit-Wolf estimator [107].

We centered the connectivity matrices according to a previously described procedure that leverages the natural geometry of the space of the covariance matrices [56–58]. First, a grand mean covariance matrix, 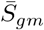, was computed by taking the geometric mean covariance matrix across all participants and epochs. Then, for each participant *i* we computed the geometric mean covariance matrix across task epochs, 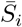, and each task epoch covariance matrix *S*_*ij*_ was projected onto the tangent space at this mean participant covariance matrix 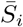 to obtain tangent vector

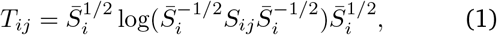

where log denotes the matrix logarithm. We then transported each tangent vector to the grand mean 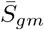 using the transport proposed by [108], obtaining a centered tangent vector

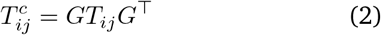

where 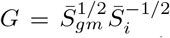. Finally, we projected each centered tangent vector back onto the space of covariance matrices, to obtain the centered covariance matrix

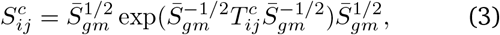

where exp denotes the matrix exponential. For the benefits (and general necessity) of this centering approach, see Fig. 1D, and for an additional overview, see [57].

### Manifold construction

Connectivity manifolds for all centered functional connectivity matrices were derived in the following steps. First, consistent with previous studies [38, 47], we applied row-wise thresholding to retain the top 10% connections in each row, and then computed cosine similarity between each row to produce an affinity matrix that describes the similarity of each region’s connectivity profiles. Second, we applied principal components analysis (PCA) to obtain a set of principal components (PC) that provide a low-dimensional representation of connectivity structure (i.e. connectivity gradients). We selected PCA as our dimension reduction technique based on recent work demonstrating the improved reliability of PCA over non-linear dimensionality reduction techniques (e.g., diffusion map embedding; [61]).

To provide a basis for comparing changes in functional network architecture that arise during learning specifically, we constructed a template manifold from a group-average Baseline connectivity matrix derived from the geometric mean (across participants) of all centered Baseline connectivity matrices. This template Baseline manifold, which underwent the same aforementioned manifold construction procedures, served as a representative task Baseline space, and as such, we aligned all individual manifolds (32 participants × 4 epochs; 128 total) to the template manifold using Procrustes alignment. All analyses on the aligned manifolds were performed using the top three PCs, which cumulatively explained 49.30% of the total variance in the template manifold. Although PC4 (8.98%) explained a similar amount of variance to PC3 (9.63%), including PC4 did not substantially impact the results and interpretations of our main analyses (see Supplementary Fig. S1). Across participants and epochs, the top three PCs, following Procrustes alignment, had an average correlation of *r* = .89 with their respective PCs in the template manifold, thus demonstrating good overall reliability and alignment across participants and epochs. Together, this approach enabled us to selectively examine adaptation-related changes in low-dimensional connectivity structure with respect to a well-defined Baseline task functional architecture, thus improving the sensitivity of our analyses.

### Manifold eccentricity

Recent work has quantified the embedding of regions and networks in low-dimensional spaces using Euclidean distance [50, 51, 67, 109]. Eccentricity refers to the Euclidean distance between a single region and the manifold centroid [51], which is equivalent to a region’s magnitude, or vector length, in the case of PCA. Eccentricity provides a scalar index of network integration and segregation, in which distal regions with greater eccentricity are more segregated than proximal regions that integrate more broadly across functional networks [51, 67, 109]. To validate this interpretation with our own data, we correlated template manifold eccentricity with three graph theoretical measures of functional integration and segregation. These measures were calculated on the row-wise thresholded template connectivity matrix and included node strength, which is the sum of a region’s connectivity weights; within-module degree *z*-score, which measures the degree centrality of a region within its respective network; and participation coefficient, which measures the network diversity of a region’s connectivity distribution [110]. Regions were assigned to their respective intrinsic functional networks [54, 62] for within-module degree *z*-score and participation coefficient. The significance of each correlation was evaluated against a null distribution generated by projecting each measure (node strength, within-module degree *z*-score, or participation coefficient) onto their respective Schaefer 1000 parcels on the 32k *fsLR* spherical mesh and performing 1000 iterations of the Cornblath spin permutation procedure [111, 112].

We computed eccentricity for each region for all individual manifolds (each participant and epoch). This allowed us to observe manifold expansions (increases in eccentricity) and contractions (decreases in eccentricity) throughout adaptation, thereby probing adaptation-related changes in functional integration and segregation.

### Eccentricity analyses

We compared region eccentricity between Baseline, Early and Late adaptation epochs by performing repeated-measures ANOVAs for each region. Results were corrected for multiple comparisons using false discovery-rate (FDR) correction at *q <* .05 [113]. We performed additional *post hoc* paired *t*-tests on significant regions to identify significant changes in eccentricity between individual epochs; FDR correction was applied (*q <* .05) across all comparisons (998 regions × 3 contrasts).

To succinctly describe and interpret the results of our region-wise analyses, we performed *k*-means clustering on regions exhibiting significant adaptation-related effects. As adjacent regions in low-dimensional space during Baseline tend to give rise to similar effects during adaptation (see Fig. 3C), clustering was performed using each region’s average embedding during Baseline (i.e. the average loading for each of the three PCs), therefore identifying ensembles of regions with common effects. *k* = 4 provided a parsimonious solution reflecting the four broad trends of changes observed throughout adaptation (see Fig. 3C). As the purpose of this analysis was to simply summarize our region-wise effects, we performed no statistical analyses directly on the different ensembles.

The primary aim of the current study was to examine changes in functional brain organization during adaptation. However, for completeness and curious readers, we provide a supplemental analysis on the subsequent de-adaptation (i.e. Washout) that unfolds as the cursor perturbation is removed (see Supplemental Fig. 4). Here, we performed planned contrasts (region-wise paired *t*-tests) between Washout and Baseline, Early, and Late adaptation. FDR correction was applied (*q <* .05) across all comparisons (998 regions × 3 contrasts). To visualize the (dis)similarity between Washout and the remaining epochs, we used UMAP [59] to derive a two-dimensional embedding based on the pairwise Euclidean distances between epochs, which was computed using the eccentricity of regions with significant adaptation-related effects (i.e. all regions in Fig. 3B). Therefore, this visualization situated de-adaptation (Washout) effects with respect to the regions that exhibited adaptation-related effects over the other phases of the task (Baseline, Early and Late adaptation). As seen in Fig. S4, de-adaptation (Washout) is associated with an overall contraction of sensorimotor and DMN areas along the cortical manifold, with Washout exhibiting closest similarity to the pattern of effects observed during Early adaptation (Fig. S4B).

### Seed connectivity analyses

In order to explore the underlying changes in functional connectivity that ultimately give way to changes in manifold eccentricity, we performed seed connectivity contrasts between the different task epochs. We selected seed regions according to the following procedure. First, to examine key connectivity differences between Early adaptation and Baseline, we selected the three largest clusters in the Early*>*Baseline contrast (Fig. 3E), which included left PMC, SPC, and PMd. Second, because regions within the same cluster tend exhibit the same overall effect (Fig. 3C-D), within each cluster we selected the region with the *t*-value closest to the cluster average, thus allowing us to demonstrate the average effect of the cluster. To probe changes in visual cortex during Late adaptation, we repeated our seed contrast analysis using a Late*>*Early contrast. A representative seed region in left extrastriate cortex was identified in the largest cluster from the Late*>*Early contrast (Fig. 3E) using the same seed selection procedures as above.

For each seed, we generated functional connectivity maps for the epochs of interest in every participant. For PMC, SSc/SPC, and PMd seeds, we performed an Early*>*Baseline seed contrast by computing region-wise paired *t*-tests. For the visual cortex seed, we performed a Late*>*Early contrast. Note that we also performed an Early*>*Baseline contrast with the visual cortex seed region, as this region also displays a significant reduction in eccentricity between Early and Baseline (see Fig. 3E, Early*>*Baseline). For all contrasts, we opted to show unthresholded *t*-maps so as to visualize the complete multivariate pattern of connectivity changes that drive changes in eccentricity. Indeed, these analyses are mainly intended to provide characterization (and interpretation) of the connectivity changes of representative regions from our main eccentricity analyses.

### Behavioural correlation analyses

We performed two correlational analyses to investigate the relationship between manifold structure and individual differences in performance during initial adaptation. First, we computed a correlation, across participants, between each region’s eccentricity during Early adaptation and the median angular error of the 120 trials completed during Early adaptation (Early error). This produced a correlation map between participants’ Early eccentricity and Early error. FDR correction (*q <* .05) was applied to correct for multiple comparisons across all regions.

Second, to explore correlations at the level of whole-brain functional networks, we took the average eccentricity, across regions, within each functional network for every participant and correlated this ‘network eccentricity’ with participant Early error. Here, we used the 17-network Schaefer 1000 assignments [54] in order to capture the spatial specificity of the region-wise correlation map, such as the difference in brain-behaviour correlations between dorsal and ventral somatomotor regions (see Fig. 5C). We applied an FDR correction (*q <* .05) across all 17 correlation tests to correct for multiple comparisons. Together, these complementary approaches enabled us to explore how individual differences in performance relate to manifold structure at both the region- and network-levels.

### Software

All code used for analyses is available on Github [https://github.com/danjgale/adaptation-manifolds]. All analyses were performed using Python 3.8.5 and involved the following open-source Python packages. Functional connectivity estimation and centering were performed with *Nilearn* 0.7.1 [92] and *PyReimann* 0.2.6 [114], respectively. All steps to generate and align connectivity manifolds were generated using *Brainspace* 0.1.1 [60]. Graph theoretical measures were computed using *Brain Connectivity Toolbox* (bctpy; https://github.com/aestrivex/bctpy/wiki), and spin permutation testing procedures were implemented in *neuromaps* 0.0.1 [115]. All statistical analyses were performed with *Pinguoin* 0.5.0 [116] and *Scipy* 1.7.2 [117]. For unsupervised learning analyses, UMAP was implemented with *Umap-learn* 0.5.2 [59], and *k*-means clustering was performed with *Scikit-learn* 0.24.1 [118]. Surface visualizations were generated using *Surfplot* [119]. All other general data processing and visualization was performed using *Numpy* 1.19.2 [120], *Pandas* 1.2.3 [121, 122], *Nibabel* 3.2.1 [123], *Matplotlib* 3.4.2 [124], *Seaborn* 0.11.1 [125], and *Cmasher* 1.6.1 [126].

---

**Fig. S1.**
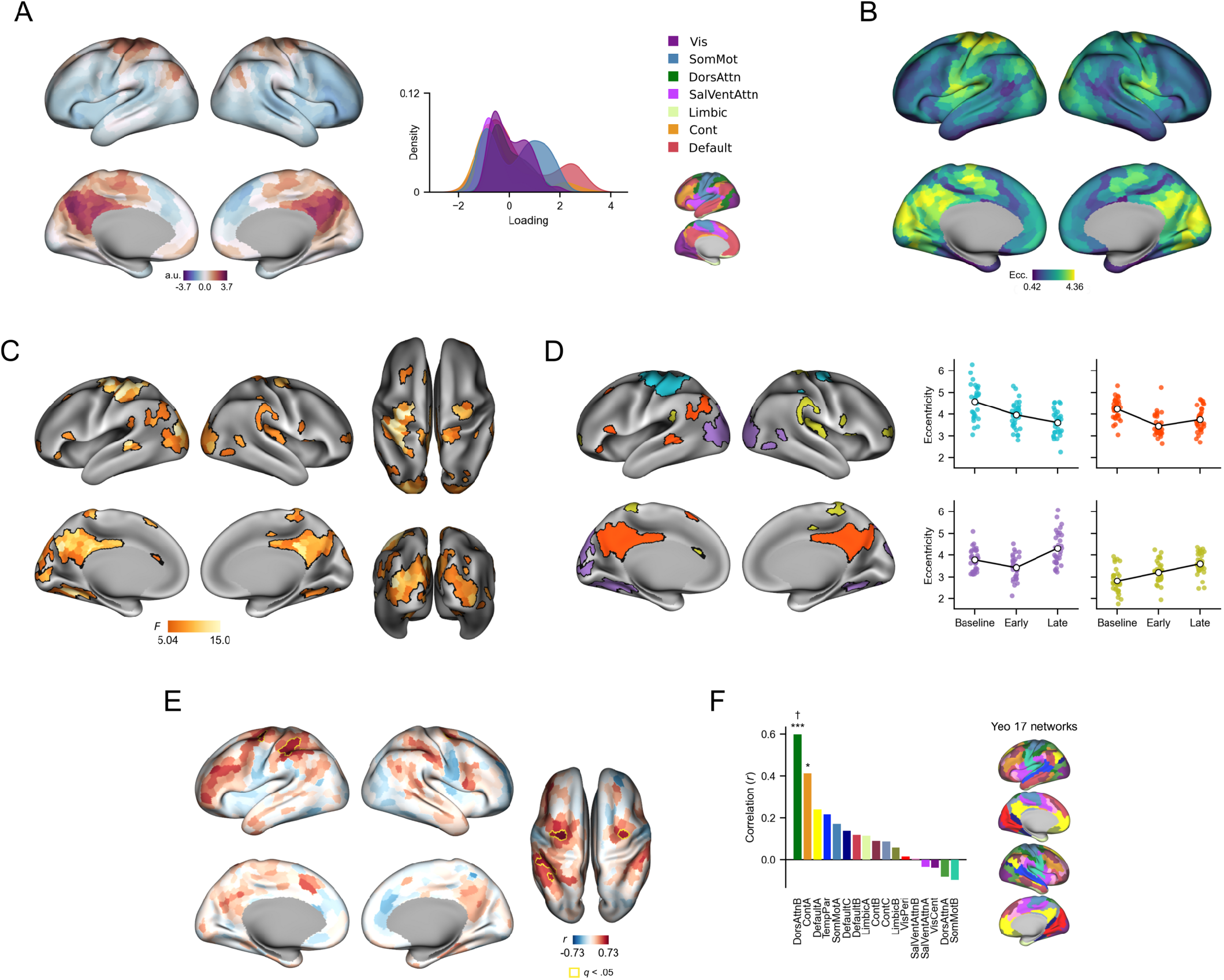
Four-dimensional manifold with the inclusion of PC 4. (*A*) Overview of PC 4. PC 4 primarily distinguishes dorsal somatomotor and PMC regions from other brain networks. Note that these regions show strong task effects (see *C*, Fig. 3*B*), which suggests that PC 4 represents a task-specific component. (*B*) Manifold eccentricity using PCs 1-4. While overall patterns of eccentricity remain similar to Fig. 2*E*, eccentricity is expectedly enhanced for dorsal somatomotor and PMC regions due to their distinction in PC 4. (*C*) Same as Fig. 3*B*, but using PCs 1-4. Overall, task effects remain similar to Fig. 3*B*, with the exception of more pronounced effects in bilateral PMC, which likely reflect more sensitivity to connectivity differences in these regions due to PC 4. (*D*) Same as Fig. 3*D*, using data from *C*. (*E*) Same as Fig. 5*C*, but using PCs 1-4. (*F*) Same as Fig. 5*E*, but using PCs 1-4. Overall, the correlations between eccentricity and performance during Early adaptation remain largely unaffected by the inclusion of PC 4.

**Fig. S2.**
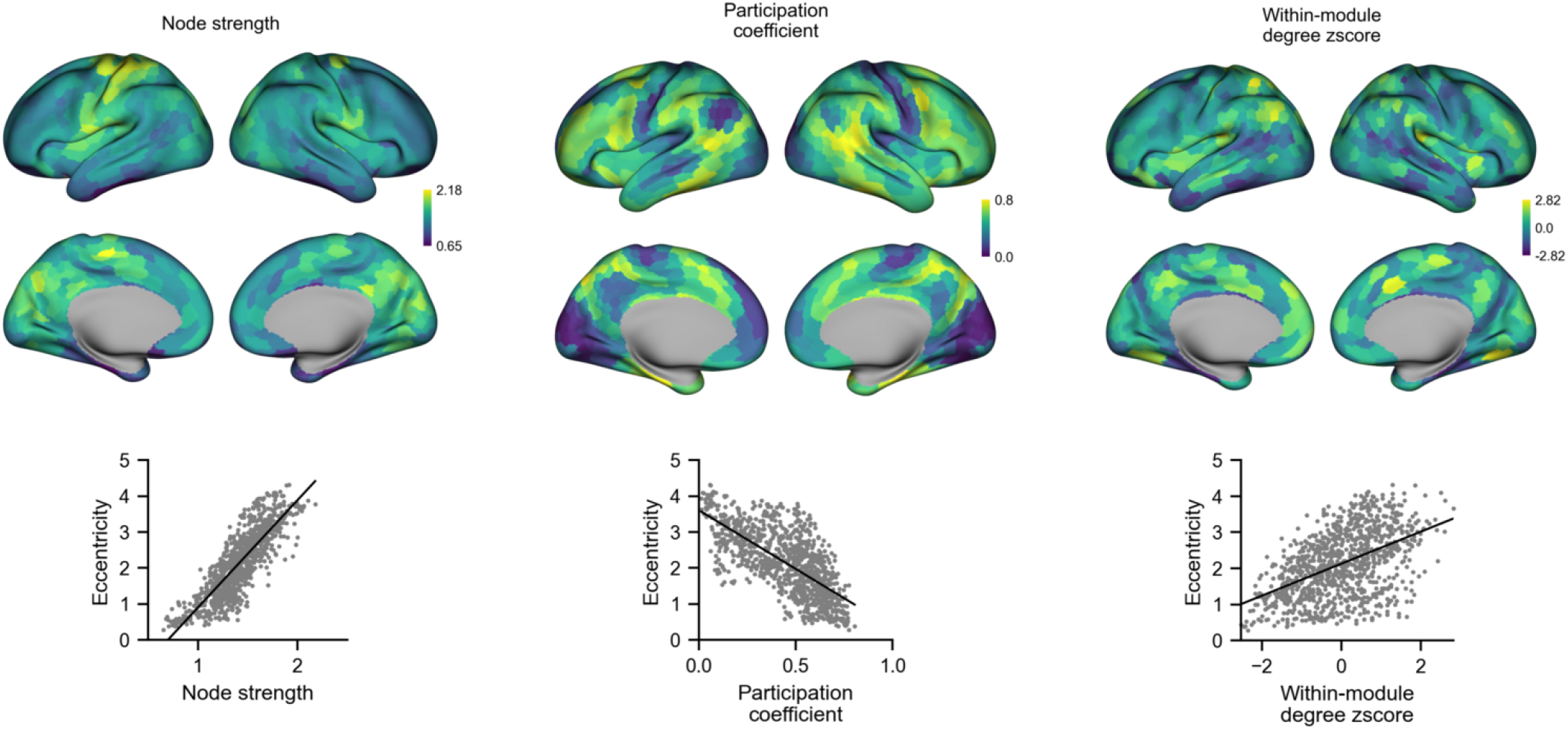
Functional connectivity properties underlying manifold eccentricity. *Top*, maps of node strength, participation coefficient, and within-module degree *z*-score from the group-average Baseline connectivity matrix (i.e. reference connectivity matrix). *Bottom*, corresponding correlations between each functional connectivity measure and manifold eccentricity (ordinary least-squares linear regression line overlaid).

**Fig. S3.**
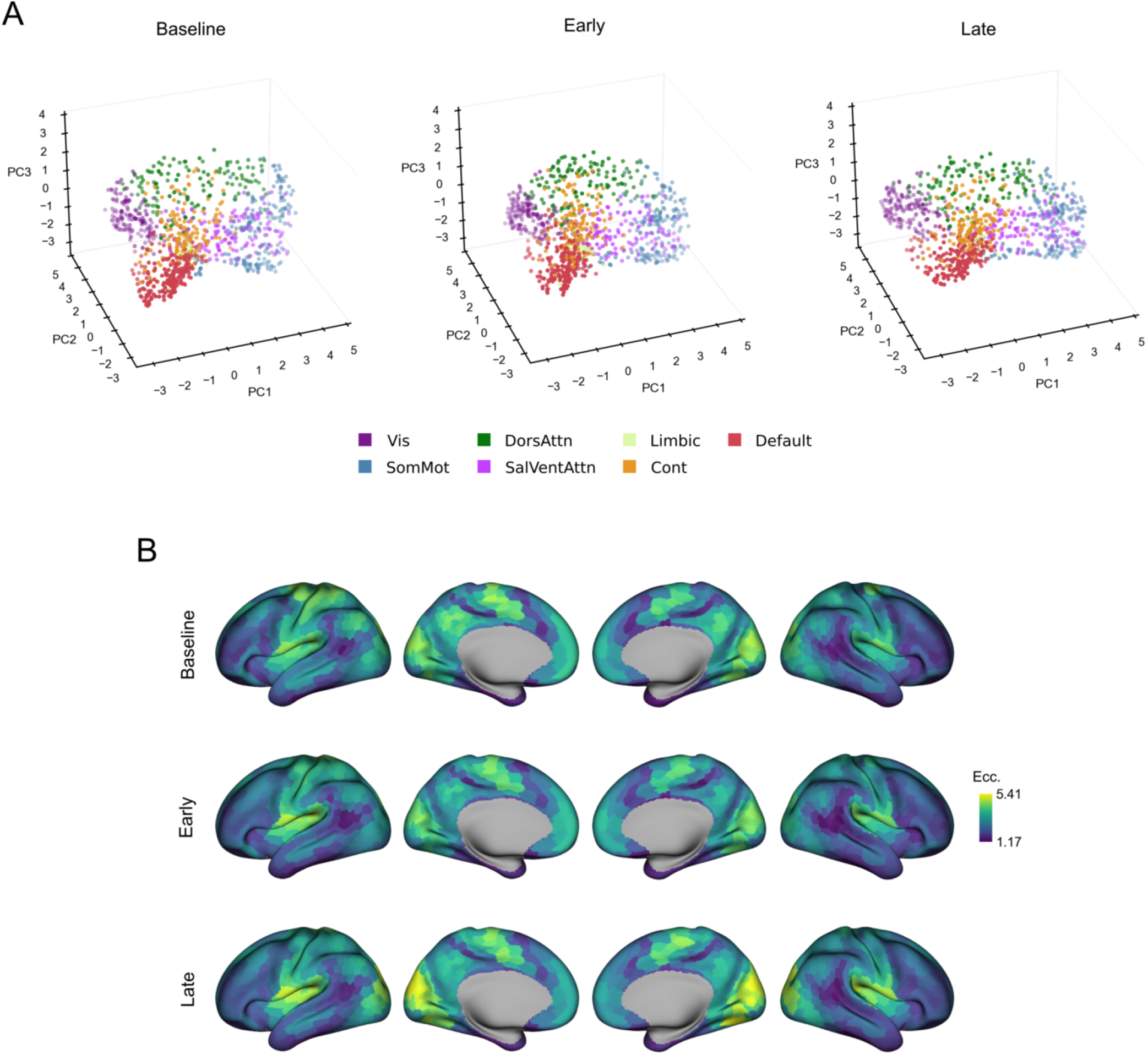
Group-average manifold structure and eccentricity across task epochs. (*A*) Mean region embeddings. (*B*) Raw mean eccentricity maps.

**Fig. S4.**
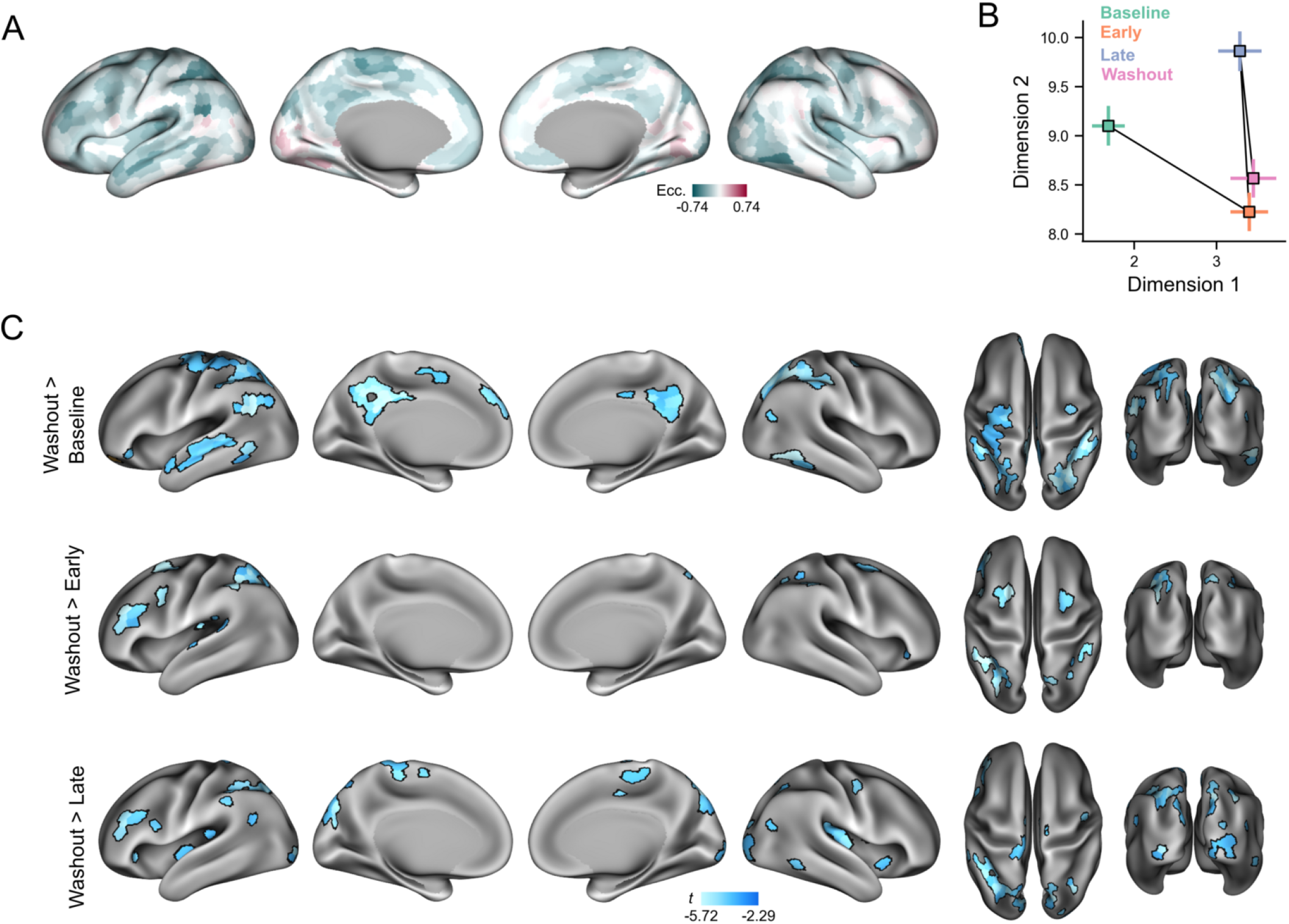
Washout eccentricity. (*A*) Average change in eccentricity during Washout relative to Baseline from Fig. 3*A*. (*B*) Two-dimensional embedding of task epochs using the UMAP algorithm [59]. To situate each epoch based on adaptation-related regions, embedding was performed on the eccentricity values of significant regions in Fig. 3*B* for each participant and epoch. Square markers indicate mean embedding across participants, and bars show ±1 SEM of each dimension. (*C*) Pairwise contrasts of eccentricity between Washout and remaining epochs. Region-wise paired *t*-tests were performed for each contrast and FDR correction (*q <* .05) was applied across all comparisons.

